# Altered skeletal muscle glucose-fatty acid flux in amyotrophic lateral sclerosis (ALS)

**DOI:** 10.1101/2020.04.02.021238

**Authors:** Frederik J. Steyn, Siobhan E. Kirk, Tesfaye W. Tefera, Teresa Y. Xie, Timothy J. Tracey, Dean Kelk, Elyse Wimberger, Fleur C. Garton, Llion Roberts, Sarah Chapman, Jeff S. Coombes, W. Matthew Leevy, Alberto Ferri, Cristiana Valle, Frédérique René, Jean-Philippe Loeffler, Pamela A. McCombe, Robert D. Henderson, Shyuan T. Ngo

**Author notes:** **To whom correspondence should be addressed:** Dr Shyuan Ngo, The Australian Institute for Bioengineering and Nanotechnology, The University of Queensland, St Lucia, Brisbane, 4072, Australia.

## Abstract

Amyotrophic lateral sclerosis (ALS) is characterized by the degeneration of upper and lower motor neurons, yet an increasing number of studies in both mouse models and patients with ALS suggest that altered metabolic homeostasis is a feature of disease. Pre-clinical and clinical studies have shown that modulation of energy balance can be beneficial in ALS. However, our capacity to target specific metabolic pathways or mechanisms requires detailed understanding of metabolic dysregulation in ALS. Here, using the SOD1^G93A^ mouse model of ALS, we demonstrate that an increase in whole-body metabolism occurs at a time when glycolytic muscle exhibits an increased dependence on fatty acid oxidation. Using myotubes derived from muscle of ALS patients, we also show that increased dependence on fatty acid oxidation is associated with increased whole-body energy expenditure. In the present study, increased fatty acid oxidation was associated with slower disease progression. However, we observed considerable heterogeneity in whole-body metabolism and fuel oxidation profiles across our patient cohort. Thus, future studies that decipher specific metabolic changes at an individual patient level are essential for the development of treatments that aim to target metabolic pathways in ALS.

## Introduction

Amyotrophic lateral sclerosis (ALS) is a fatal neurodegenerative disease that is characterized by the degeneration of motor neurons in the brain, brainstem and spinal cord. The progressive loss of neurons in ALS results in muscle denervation, weakness and paralysis. Death usually occurs within 3-5 years from diagnosis (Brown and Al-Chalabi, 2017). ALS is a highly variable disease in terms of age of onset, site of symptom onset, and rate and pattern of disease progression. Superimposed on the clinical variability of ALS is the underlying complexity of genetic contribution and the pathogenic pathways that lead to the death of neurons (Brown and Al-Chalabi, 2017; Ghasemi and Brown, 2017).

While multiple pathogenic mechanisms, including neuronal hyperexcitability (Vucic *et al*., 2008; Kiernan, 2009), glutamate excitotoxicity (Rothstein *et al*., 1992), protein aggregation (Arai *et al*., 2006), and oxidative stress (Blasco *et al*., 2017), are proposed to underpin disease pathology, there are an increasing number of studies that emphasize a negative role of metabolic dysfunction on disease progression (Dupuis *et al*., 2004; Ahmed *et al*., 2016; Ioannides *et al*., 2016). In human ALS, an increase in resting energy expenditure (i.e. hypermetabolism) has been observed in approximately one third to more than half of all patients (Desport *et al*., 2001; Desport *et al*., 2005; Bouteloup *et al*., 2009; Funalot *et al*., 2009; Vaisman *et al*., 2009; Jesus *et al*., 2018; Steyn *et al*., 2018a). More recently, we have shown that hypermetabolism in patients with ALS is associated with greater clinical evidence of lower motor neuron dysfunction, more aggressive disease progression, and increased risk of earlier death (Steyn *et al*., 2018a). Despite this clear evidence of the importance of metabolism, the capacity to target metabolism to slow disease progression is restricted by our limited understanding of the underlying biochemistry of metabolism in ALS.

Previous studies in SOD1^G93A^ and SOD1^G86R^ mouse models of ALS suggest that altered metabolic balance is a feature of the disease (Dupuis *et al*., 2004), with early and persistent perturbations in the blood levels of metabolic hormones such as leptin, the presence of circulating ketone bodies, and increased energy expenditure (Dupuis *et al*., 2004). Reports of increased peripheral clearance of circulating lipids (Fergani *et al*., 2007), and increased expression of fatty acid metabolism genes in glycolytic muscle (Palamiuc *et al*., 2015) point towards altered skeletal muscle metabolism as a potential contributor to the metabolic phenotypes observed in ALS. However, whether this altered profile of gene expression translates into functional changes in glucose-fatty acid metabolism in the muscle of ALS mice, and how this might relate to metabolic changes at the whole-body level remains to be determined. Moreover, whether similar metabolic changes occur in the skeletal muscle of ALS patients, and the relevance of these changes to the clinical and metabolic presentations in human ALS is unknown.

In this study, we aimed to address this gap in knowledge by characterizing whole-body metabolism in the SOD1^G93A^ mouse model of ALS throughout the course of disease, and by assessing energy substrate utilization in intact glycolytic muscle fibers isolated from SOD1^G93A^ mice when compared to wild-type (WT) controls. We also assessed fatty acid and glucose oxidation in myotubes derived from well characterized ALS patients and healthy controls to determine whether substrate metabolism in ALS muscle is linked to the energy expenditure profiles and clinical features. We report evidence of a functional alteration in glucose-fatty flux in glycolytic muscle of SOD1^G93A^ mice. In ALS patient-derived myotubes, changes in glucose-fatty acid oxidation are associated with whole-body energy expenditure and disease progression, but not hypermetabolism.

## Methods

### Animal studies

#### Mice

Experiments at the University of Queensland were approved by the University of Queensland Animal Ethics Committee and conducted in accordance with the Queensland Government Animal Care and Protection Act 2001, associated Animal Care and Protection Regulations (2002 and 2008), and the Australian Code of Practice for the Care and Use of Animals for Scientific Purposes (2004). Experiments at the University of Notre Dame were conducted in accordance with the FLSC Guidelines and the Notre Dame IACUC Policy on Humane Endpoints for animal use.

Transgenic mice overexpressing the human *SOD1* G93A mutation (B6-Cg-Tg (SOD1-G93A) 1Gur/J) (Gurney *et al*., 1994) were purchased from the Jackson Laboratory (Bar Harbor, ME, USA) and bred on a C57BL/6J background. Male SOD1^G93A^ mice and litter- or age-matched WT control mice were used for experiments at ages that correspond to pre-defined stages of disease in SOD1^G93A^ mice; presymptomatic (5 weeks of age, no symptoms), onset (9-11 weeks of age, early signs of hindlimb tremor and weakness), mid-stage (16-19 weeks of age, pronounced hindlimb weakness), and end-stage (21-25 weeks of age, significant hindlimb weakness leading to paralysis and euthanasia due to loss of the righting reflex) (Ngo *et al*., 2012; Lee *et al*., 2013). Mice were randomly assigned to experiments and matched by litter, or by age. All mice were group-housed (3-4 mice per cage) in filter top cages or in indivudally ventilated cages (IVC) when maintained in a specific pathogen free environment. 12h light, 12h dark cycle (on at 0600h and off at 1800h) and had free access to food (20% protein, 4.8% fat; Specialty Feeds, WA, AUS) and water. Room temperature was maintained at 22 ± 2°C. Prior to the tissue collection, all mice were anesthetized with an intraperitoneal injection of sodium pentobarbitone (32.5 mg/kg, Virbac Animal Health, NSW, AUS). Following complete loss of the pedal withdrawal reflex and eye-blink reflex, mice were killed by cervical dislocation. All animal work was conducted in accordance with the ARRIVE guidelines (Kilkenny *et al*., 2010).

### EchoMRI assessment of fat and fat free mass

All imaging took place in the Notre Dame Integrated Imaging Facility Friemann Life Sciences Center. Whole body fat and fat free mass was measured with an EchoMRI-130™ QMR (EchoMRI, TX, USA) (Metzinger *et al*., 2014). Two X-ray images were taken at different energy levels to assess soft tissue density and bone. Mass measurements for fat and fat free tissue were produced for all scans in the EchoMRI Body Composition Analyzer EMR-184 software (EchoMRI).

### Indirect calorimetry

Energy expenditure was measured with a Phenomaster open-circuit indirect calorimetry system housed within a temperature (22°C) and 12h light, 12h dark cycle (on at 0600h and off at 1800h) controlled chamber (TSE-Systems, Bad Homburg, DEU), as we have done previously (Steyn *et al*., 2018c). Experimental cages (n=16) were sampled at 60min intervals for 3.5 min/cage, with concentrations of O_2_ and CO_2_ in the outgoing air being measured sequentially within each interval. One vacant cage was included to obtain a reference concentration for ambient gas. Activity (x- and y-plane), food intake, and body weight was recorded synchronously with metabolic data. Measurements were performed continuously over 72h, with analysis restricted to the final 24h assessment period (allowing 48h of acclimation). For data analysis, measures of total energy expenditure and food intake were adjusted for body weight.

### ^18^F-deoxyglucose (FDG) PET/Single-photon emission computed tomography (SPECT)/CT imaging

Animals were anesthetized using isoflurane inhalation for 2-5min in a vaporized-controlled tank. Animal breathing was checked once every minute by visual inspection. ^18^F-deoxyglucose (FDG) at a dose of 0.200mCI activity was administered to mice intravenously through tail vein (volume <100μl). PET images were acquired on a trimodal Alibra PET/SPECT/CT image station (Carestream Health, Woodbridge, CT) to produce high-resolution PET images that were reconstructed for analysis. FDG uptake in brown adipose tissue was quantified as the mean voxel value within a visually determined volume of interest as described previously (van der Veen *et al*., 2012).

### Ex vivo lipolysis

Epididymal white adipose tissue (WAT) was excised and rinsed in 1 x PBS supplemented with 0.1% fatty acid-free BSA (FAF-BSA). All epididymal WAT explants were placed in plastic vials containing 1mL of modified Kreb’s-Henseleit buffer (in mM): 4.7 KCL, 1.2 KH_2_PO_4_, 1.2 MgSO_4_.7H_2_O, 1.25 CaCl_2_.2H_2_O, 25 NaHCO3, 5 Glucose, 118 NaCl and 4% FAF-BSA. Buffer was gassed with 95% O_2_/5% CO_2_ for 45min to reach a pH of 7.4. All procedures were conducted in a shaking water bath at 37°C. For the assessment of non-esterified fatty acids (NEFA), 6μl of buffer was collected at 0, 30, 60, 90 and 120min time points, placed immediately on dry ice, and stored at −80°C. Samples were assayed on a NEFA-C kit (Wako Chemicals, Osaka, JPN). For glycerol, buffer was collected after 2h of incubation. Glycerol content was assessed using a free glycerol determination kit with glycerol standard solution (Sigma-Aldrich, MO, USA). Final glycerol content and NEFA-C levels were expressed relative to the weight of the respective epididymal WAT explant.

### Plasma NEFA

Following sacrifice, terminal blood samples were collected from mice via cardiac puncture. Samples were transferred into EDTA-precoated tubes and centrifuged for 3min. Plasma was aliquoted and stored at −80°C until use. NEFA levels in plasma were determined using a NEFA C-test kit (Wako Chemicals).

### Oil Red O staining

Extensor digitorum longus (EDL) muscles were embedded in optimum cutting temperature compound and rapidly frozen in liquid nitrogen cooled isopentane. Oil Red O (ORO) staining was performed on muscle cryosections (10μm) to visualize neutral lipids using an ORO kit (Abcam, Cambridge, GBR). Briefly, sections were fixed with 10% neutral buffered formalin for 15min and rinsed three times with distilled water for 30s at room temperature. ORO solution was added onto the sections and incubated for 10min, and the slides were differentiated in 85% propylene glycol solution for 1min at room temperature. After two rinses, slides were air dried and mounted with aqueous mounting agent (Aquatex, EMD Millipore, CA, USA). Bright-field images (20× magnification) were taken with an Aperio ScanScope system (Leica, Mannheim, DEU). ORO labeling was quantified using 10+ randomly selected sections of muscle per animal (n=5 per group). Representative images of the muscle section were processed using ImageJ to identify the mean ORO intensity for each animal, following thresholding to remove non-specific labeling (threshold set to 225 (0-255)).

### Assessment of cellular respiration in muscle fiber bundles

EDL muscle fiber bundles were chemically dissociated as previously described (Li *et al*., 2016). Muscle fiber bundles were seeded onto Seahorse XF^e^96 microplates in culture media (low glucose DMEM supplemented with 10% FBS and 1% Antibiotic-Antimycotic; ThermoFisher, MA, USA), and maintained overnight at 37°C with 5% CO_2_. Prior to the commencement of metabolic assays, muscle fiber viability was assessed using an alamarBlue cell viability assay (ThermoFisher) and used for the data normalization. Real-time assessment of bioenergetic parameters in EDL fiber bundles was performed on the XF^e^96 Extracellular Flux Analyzer (Agilent Technologies, CA, USA).

The dependence and capacity of EDL fibers to use glucose and fatty acid as fuel substrates was determined using the Seahorse XF Mito Fuel Flex Test Kit (Agilent Technologies). Prior to the assay, culture media was replaced with pre-warmed assay media (pH 7.4) consisting of XF base media (Agilent Technologies), 10mM D-glucose (Sigma-Aldrich), 1mM sodium pyruvate (Sigma-Aldrich) and 2mM L-glutamine (ThermoFisher). Carnitine palmitoyltransferase 1A inhibitor etomoxir (ETO, 4.0μM), mitochondrial pyruvate carrier inhibitor UK5099 (2.0μM) and glutaminase inhibitor BPTES (3.0μM) were prepared with assay media and loaded into the XF^e^96 sensor cartridge following the manufacturer guidelines. Following the first 3 cycles of baseline measurement of oxygen consumption rate (OCR), the decrease of OCR levels upon inhibition of one or more pathways was continuously recorded for the following 6 cycles. Each cycle consisted of 3min mix, 30s wait and 3min measurement. The dependence and capacity of each fuel pathway relative to total fuel oxidation was calculated according to the manufacturer guidelines.

Substrate induced maximal respiration was tested in EDL fiber bundles as previously described (Li *et al*., 2016). Briefly, plates containing muscles fibers were changed into assay media (pH 7.4) containing 120mM NaCl, 3.5mM KCl, 1.3mM CaCl_2_, 0.4mM KH_2_PO_4_, 1mM MgCl_2_, 2.5mM D-glucose and 0.5mM L-carnitine prior to the assay run. OCR was continuously measured for 6 cycles after sequential injections of sodium pyruvate (10mM) or palmitate-BSA (100μM) with carbonyl cyanide-p-trifluoromethoxyphenylhydrazone (FCCP, 0.4μM), followed with antimycin/rotenone (1μM). Quantitation of Seahorse data were conducted over 8-24 technical replicates for n= 5-12 animals per group.

### Patient Studies

#### Subjects

Eighteen ALS patients who met the revised El-Escorial criteria for ALS (Brooks *et al*., 2000) were enrolled from the Royal Brisbane and Women’s Hospital (RBWH) ALS clinic for the collection of skeletal muscle biopsies. Eleven healthy control participants were also enrolled. These control individuals were the spouses, friends or family members of ALS participants. For all participants, exclusion criteria were history of a metabolic condition (e.g. Hashimoto’s disease) and diabetes mellitus. Participant details are shown in Table 1 and Supplementary Table 1. For ALS patients, the ALS Functional Rating Scale-Revised (ALSFRS-R) score, King’s stage, and ΔFRS were obtained from clinical records. All participants provided written informed consent; participant consent was obtained according to the Declaration of Helsinki. Work performed in this study was approved by the RBWH and University of Queensland human research ethics committees.

**Table 1:** Characteristics of ALS patients and healthy controls at the time of muscle biopsy collection

|  | Control<br>(n=11) | ALS<br>(n=18) | *P* |
| --- | --- | --- | --- |
| **Demographics** |  |  |  |
| Age (years) | 58.9±10.1 | 54.9±7.2 | 0.214 |
| Sex, female | 3 (27.27) | 4 (22.2) | 0.758 |
| **Anthropometric and metabolic measures** |  |  |  |
| BMI (kg/m<sup>2</sup>) | 25.9±2.7 | 26.6±4.2 | 0.644 |
| Fat Free Mass (kg) | 53.4±10.5 | 54.7±10.8 | 0.745 |
| Fat mass (%) | 31.1±9.4 | 32.4±11.5 | 0.759 |
| EEkc (kcal/day) | 1596±331.1 | 1809±336.2 | 0.107 |
| MI (%) | 108.4±15.41 | 119.5±9.6 | 0.025 |
| **Clinical scores** |  |  |  |
| Time since onset |  | 29.2±21.5 |  |
| ALSFRS-R |  | 36.7±6.1 |  |
| ΔFRS |  | -0.5±0.3 |  |
| King's Stage |  | 2.4±1.0 |  |
| FVC (% of predicted) |  | 88.5±22.1 |  |

Data presented as mean (SD) and n (%) for categorical data. Means were compared by independent t-test and proportions with the chi-square test; BMI - Body Mass Index, EEkc - Energy expenditure kilocalories, MI - metabolic index (measured EEkc/predicted EEkc), ALSFRS-R - ALS Functional Rating Scale Revised, ΔFRS - Change in ALSFRS-R since symptom onset, FVC - Forced Vital Capacity.

### Assessment of body composition and energy expenditure

Body composition (fat mass and fat free mass) was determined by whole body air displacement plethysmography using the BodPod system (Cosmed USA, Rome, ITA). Values of fat mass and fat free mass were used to predict resting energy expenditure (Steyn *et al*., 2018a). Resting energy expenditure in ALS and control participants was then measured by indirect calorimetry using a Quark RM respirometer (Cosmed) as per our established methodology (Steyn *et al*., 2018a). Controls were matched to patients with ALS by age, sex, weight, BMI and body composition. The metabolic index of each individual was derived by calculating measured resting energy expenditure as a percentage of predicted resting energy expenditure. A metabolic index of ⩾120% was defined as hypermetabolism (Steyn *et al*., 2018a).

### Muscle biopsy and culture

Muscle biopsies were collected from the vastus lateralis of one leg. Lignocaine (1%; 5ml) was injected to anaesthetize the skin and underlying fat and muscle tissue. A 10mm incision was made and advanced through the fascia of the muscle. A ~200mg sample of muscle was collected using a sterile 6mm hollow Bergstrom biopsy needle (Pelomi, Albertslund, Denmark) modified for suction (Tarnopolsky *et al*., 2011) and placed in holding media (DMEM/F12 with 0.5% gentamicin; ThermoFisher).

Primary myoblasts were isolated and cultured using a modified muscle tissue explant method (Tarnopolsky *et al*., 2011) and frozen as low passage cell stocks. Primary myogenic cells were maintained in DMEM/F12 medium supplemented with 20% FBS, 10% AmnioMAX C-100 and 0.5% gentamicin (ThermoFisher). Culture media was changed every second day and cells were passaged when they reached ~70% confluence. For experiments, primary myogenic cells were seeded at a density of 15000 cells/well into Seahorse XF^e^96 cell culture microplates. At 80% confluence, cells were differentiated into myotubes by replacing maintenance media with differentiation media consisting of DMEM/F12 medium with 2% non-inactivated horse serum and 0.5% gentamicin (ThermoFisher). Fresh differentiation medium was fed every 2 days until mature multinucleated myotubes formed. Primary myotubes underwent assessment of key parameters of glucose and fatty acid oxidation dependency and capacity using the Seahorse Mito Fuel Flex Test Kit (Agilent Technologies).

### Statistical analysis

Statistical differences were assessed using Prism 8.0a. (GraphPad Software Inc., La Jolla, CA, USA). Statistical comparisons, unless otherwise indicated, were performed with unpaired Student’s t-test, non-parametric t-test, or two-way ANOVA followed by Bonferroni multiple comparison. Linear relationships were assessed by Pearson correlation. All graphical data are presented as mean±SEM or as a Pearson correlation. Values of *P*<0.05 were considered to be statistically significant.

## Results

### Disease progression in SOD1^G93A^ mice is associated with a decline in both fat free mass and fat mass

To determine the impact of disease progression on body composition and total body weight in SOD1^G93A^ mice, we conducted a serial assessment of body weight, and whole-body fat free mass and fat by EchoMRI. Total body weight in SOD1^G93A^ mice was significantly lower than that of litter-matched WT controls by 19 weeks of age (Figure 1A). Loss of body weight was due, in part, to a loss in total fat free mass (Figure 1B). Tibialis anterior (TA) and gastrocnemius (Gastroc) weight was reduced by disease onset (Figure 1D and E), and extensor digitorum longus (EDL) mass was reduced by the mid-stage of disease (Figure 1F). When compared to age-matched WT controls, fat accumulation in SOD1^G93A^ mice slowed following disease onset, and whole-body fat mass was significantly lower from 15 weeks of age (Figure 1C). Epididymal and inguinal fat mass was lower in SOD1^G93A^ mice from the mid-stage of disease (Figure 1G and H). Consistent with the slowing in the accumulation of, and the eventual decline in adipose mass, circulating levels of leptin did not increase in SOD1^G93A^ mice throughout the disease course while the WT controls showed increased levels of leptin with age (Figure 1I).

**Figure 1:**
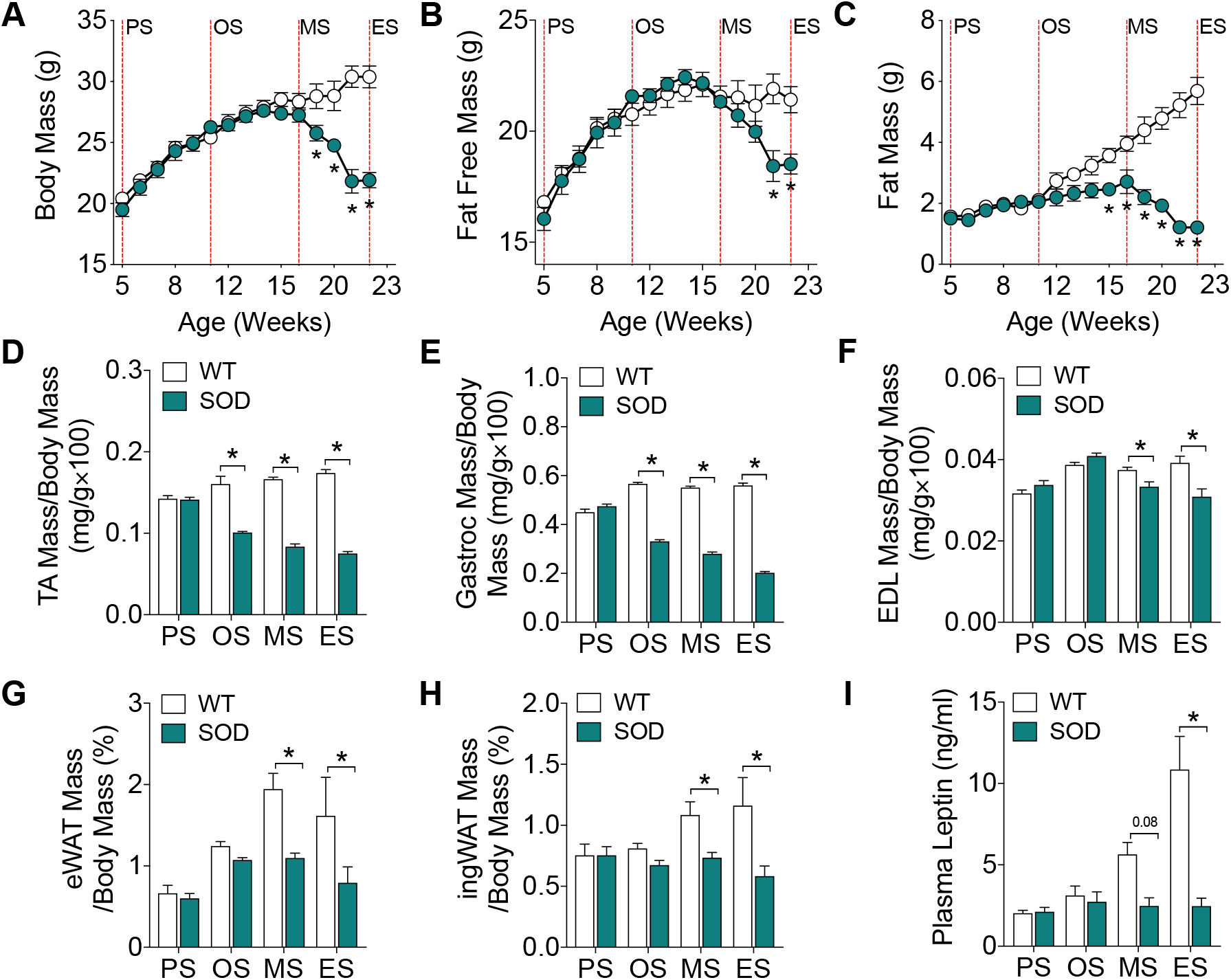
Body weight, fat free mass, and fat mass decreases in SOD1^G93A^ mice over the course of disease. (A) Total body weight (B) fat free mass and (C) fat mass in SOD1^G93A^ mice and wild-type (WT) age-matched controls (n=9-10/group). Total (D) tibialis anterior (TA), (E) gastrocnemius (Gastroc), and (F) extensor digitorum longus (EDL) muscle mass in SOD1^G93A^ mice and age-matched WT controls (n=10-26/group). Total weight of (G) epididymal white adipose tissue (eWAT) and (H) inguinal white adipose tissue (ingWAT) in SOD1^G93A^ mice and age-matched WT controls (n=5-12/group). (I) Circulating levels of leptin in SOD1^G93A^ mice and age-matched WT controls (n=5-6/group). White circles and columns represent WT mice; blue circles and columns represent SOD1^G93A^ transgenic mice. All data presented as mean ± SEM. \**P*<0.05, two-way ANOVA with Bonferroni’s post-doc test. PS, presymptomatic; OS, onset; MS, mid-stage; ES, end-stage.

### Total energy expenditure in SOD1^G93A^ mice increases by the mid-stage of disease

We next sought to determine whether energy expenditure in SOD1^G93A^ mice increases relative to disease progression. When compared to WT controls, SOD1^G93A^ mice at the mid-stage of disease had higher levels of energy expenditure during the light and dark cycle, resulting in an overall increase in 24h total energy expenditure (Figure 2A and B). SOD1^G93A^ mice had decreased activity by disease onset (Figure 2C and D).

**Figure 2:**
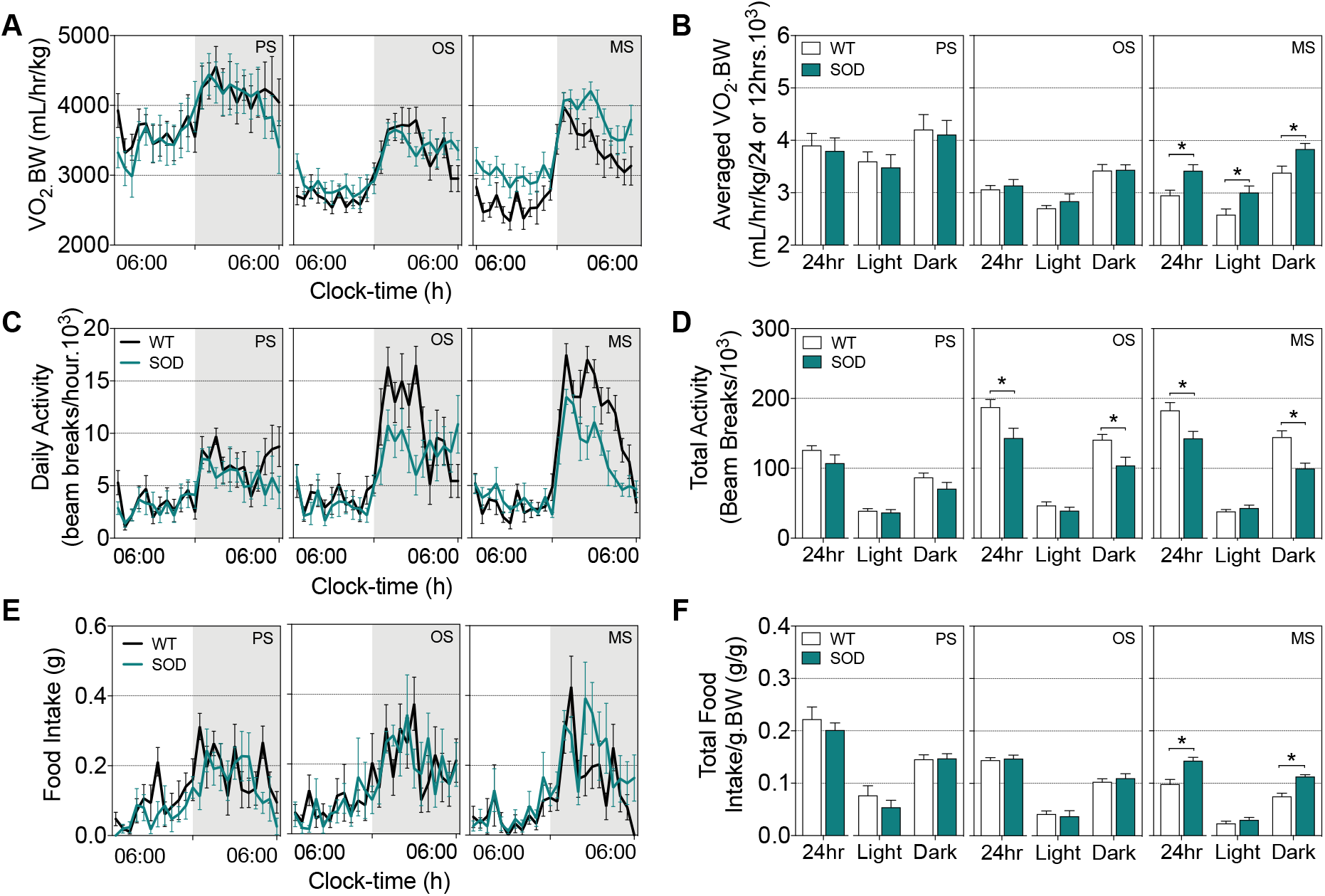
Increased energy expenditure occurs in parallel with increased food intake in symptomatic SOD1^G93A^ mice. (A) Representative data trace of oxygen consumption over 24h in SOD1^G93A^ mice and wild-type (WT) litter-matched controls. (B) Average VO_2_ consumption in these mice during the light and dark cycle, and over a 24h period. (C) Representative data trace of daily activity over 24h. (D) Average total activity during the light and dark cycle, and over a 24h period in SOD1^G93A^ mice and WT litter-matched controls. (E) Representative data trace of food intake over 24h. (F) Average food intake during the light and dark cycle, and over a 24h period in SOD1^G93A^ mice and WT litter-matched controls. Black lines and white columns represent WT mice; blue lines and columns represent SOD1^G93A^ transgenic mice. Data presented as mean ± SEM for n=8/group. \**P*<0.05, two-way ANOVA with Bonferroni’s post-doc test. PS, presymptomatic; OS, onset; MS, mid-stage.

We also investigated whether reductions in body weight and fat mass in SOD1^G93A^ mice were due to a decline in food intake. We observed no change in food intake between SOD1^G93A^ and litter-matched WT controls at disease onset. SOD1^G93A^ mice at the mid-stage of disease consumed more food than litter-matched WT controls (Figure 2E and F). Thus we find that reductions in weight are not associated with decreased food consumption, and that despite an increase in food intake and a decline in activity-dependent energy expenditure, SOD1^G93A^ mice are unable to offset increased energy expenditure to prevent the depletion of energy stores.

### Lipolytic rate is maintained in SOD1^G93A^ mice

Brown adipose tissue (BAT) regulates non-shivering thermogenesis, which itself can contribute to total energy expenditure in mice (Even and Blais, 2016). We found no difference in BAT weight (Figure 3A) or glucose uptake in BAT between SOD1^G93A^ and WT control mice (Figure 3B). Thus, the increase in energy expenditure in SOD1^G93A^ at the mid-stage of disease is unlikely to be due to increases in non-shivering thermogenesis.

**Figure 3:**
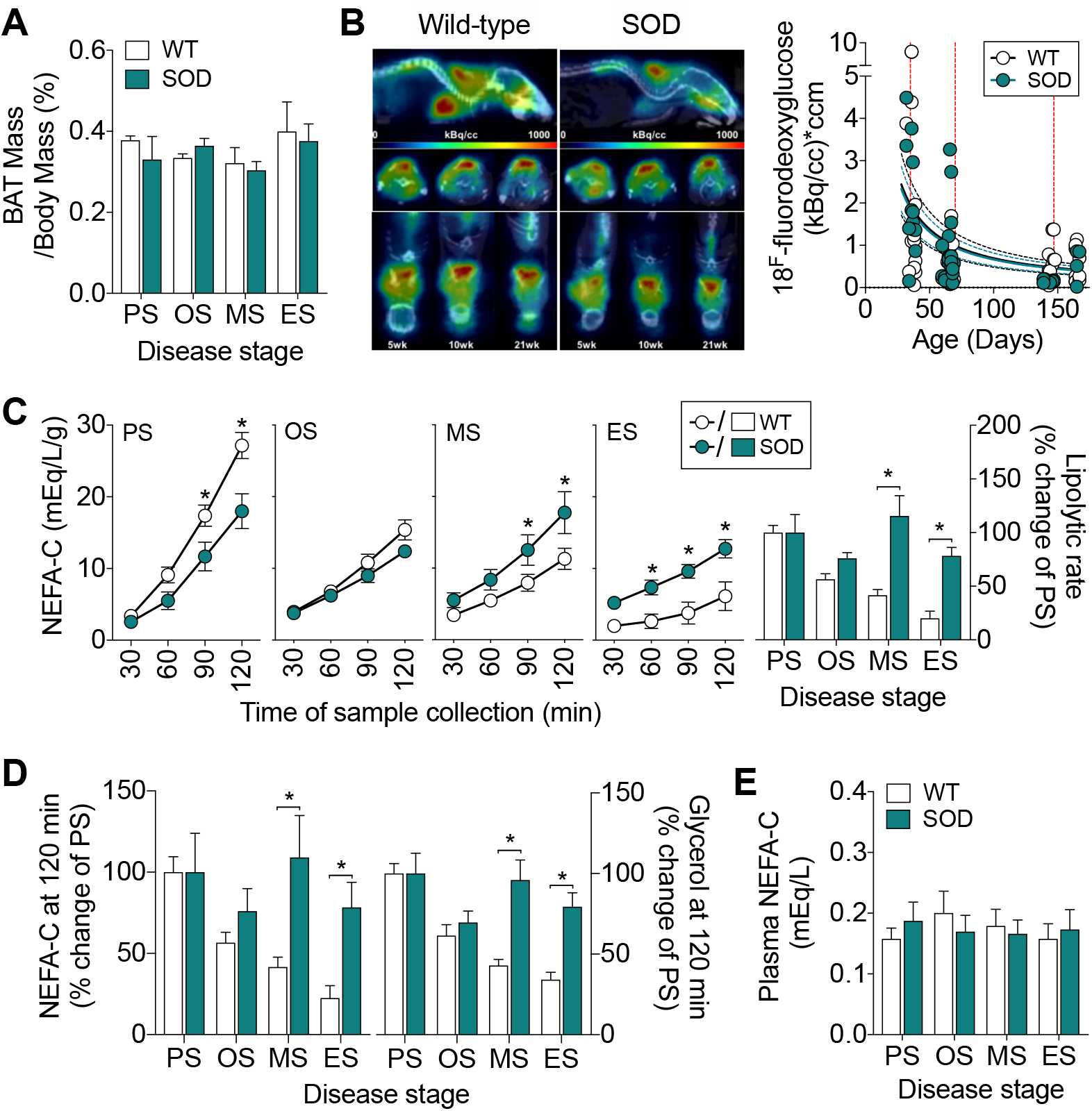
Lipolysis is maintained throughout disease course in SOD1^G93A^ mice. (A) Brown adipose tissue (BAT) weight (n=5-12/group) and (B) glucose uptake in BAT in SOD1^G93A^ mice and age-matched wild-type WT controls (n=9-11/group). (C) Levels of non-esterified fatty acids (NEFA-C) in Kreb’s-Henseleit buffer as an indicator of lipolytic rate of epididymal white adipose tissue explants from SOD1^G93A^ mice and WT controls (n=6-17/group). Lipolytic rate is expressed as a percent of that observed at PS stage. (D) Cumulative NEFA (n=6-17/group) and glycerol (n=8-17/group) in Kreb’s-Henseleit buffer. (E) Circulating plasma NEFA in SOD1^G93A^ mice and WT age-matched controls (n=12/group). White circles and columns represent WT mice; blue circles and columns represent SOD1^G93A^ transgenic mice. All data presented as mean ± SEM. \**P*<0.05, two-way ANOVA with Bonferroni’s post-doc test. PS, presymptomatic; OS, onset; MS, mid-stage; ES, end-stage.

White adipose tissue (WAT) stores lipids, providing energy reserves that are available during periods of increased energy demand. Given the slowing of accumulation and eventual reduction of fat mass in SOD1^G93A^ mice, we assessed the release of lipids (a proxy measure for lipolysis) from epididymal WAT explants. The lipolytic rate of *ex vivo* explants of epididymal WAT decreased over the lifespan of WT mice, whilst the lipolytic rate in SOD1^G93A^ mice was maintained throughout the course of disease. As such, the lipolytic rate in SOD1^G93A^ mice was significantly higher by the mid-stage of disease when compared to WT controls (Figure 3C). This corresponded to a sustained rate of appearance of cumulative non-esterified fatty acids (NEFA) and glycerol from WAT explants (Figure 3D). Circulating plasma NEFA in WT and SOD1^G93A^ mice did not differ (Figure 3E). Overall, the sustained rate of lipolysis in epididymal WAT explants from SOD1^G93A^ mice suggest that there is greater rate of turnover of lipids throughout the course of disease.

### SOD1^G93A^ mice exhibit a functional preference for fat oxidation in glycolytic EDL muscle

Previously, a decrease in the expression of glucose handling genes, and an increase in the expression of lipid handling genes in glycolytic muscle of SOD1^G86R^ mice has been suggested to drive a switch towards the use of lipid as an energy substrate (Palamiuc *et al*., 2015). Thus, sustained lipolysis in SOD1^G93A^ mice could serve to mobilize fatty acids for use as an energy source in skeletal muscle. We used oil-red O staining to quantify intramuscular lipid accumulation in the glycolytic EDL muscle, and conducted real-time assessment of substrate utilization in EDL muscle fiber bundles isolated from SOD1^G93A^ and age-matched WT mice. There was no difference in intramuscular lipid content in the EDL muscle of SOD1^G93A^ mice when compared to WT controls (Figure 4A). We observed no difference in glucose oxidation dependency (Figure 4B and C), and following the inhibition of mitochondrial fatty acid and glutamine uptake, EDL muscle fibers from SOD1^G93A^ mice exhibited similar levels of glucose oxidation capacity across all disease stages (Figure 4D and E). However, we observed increased dependence on fat oxidation in isolated EDL muscle fiber bundles from SOD1^G93A^ mice by the mid-stage of disease (Figure 4F and G). Moreover, the capacity for fat oxidation to sustain mitochondrial respiration after inhibition of pyruvate and glutamine entry into mitochondria was also increased in EDL muscle fibers bundles from SOD1^G93A^ mice at the mid-stage of disease (Figure 4H and I).

**Figure 4:**
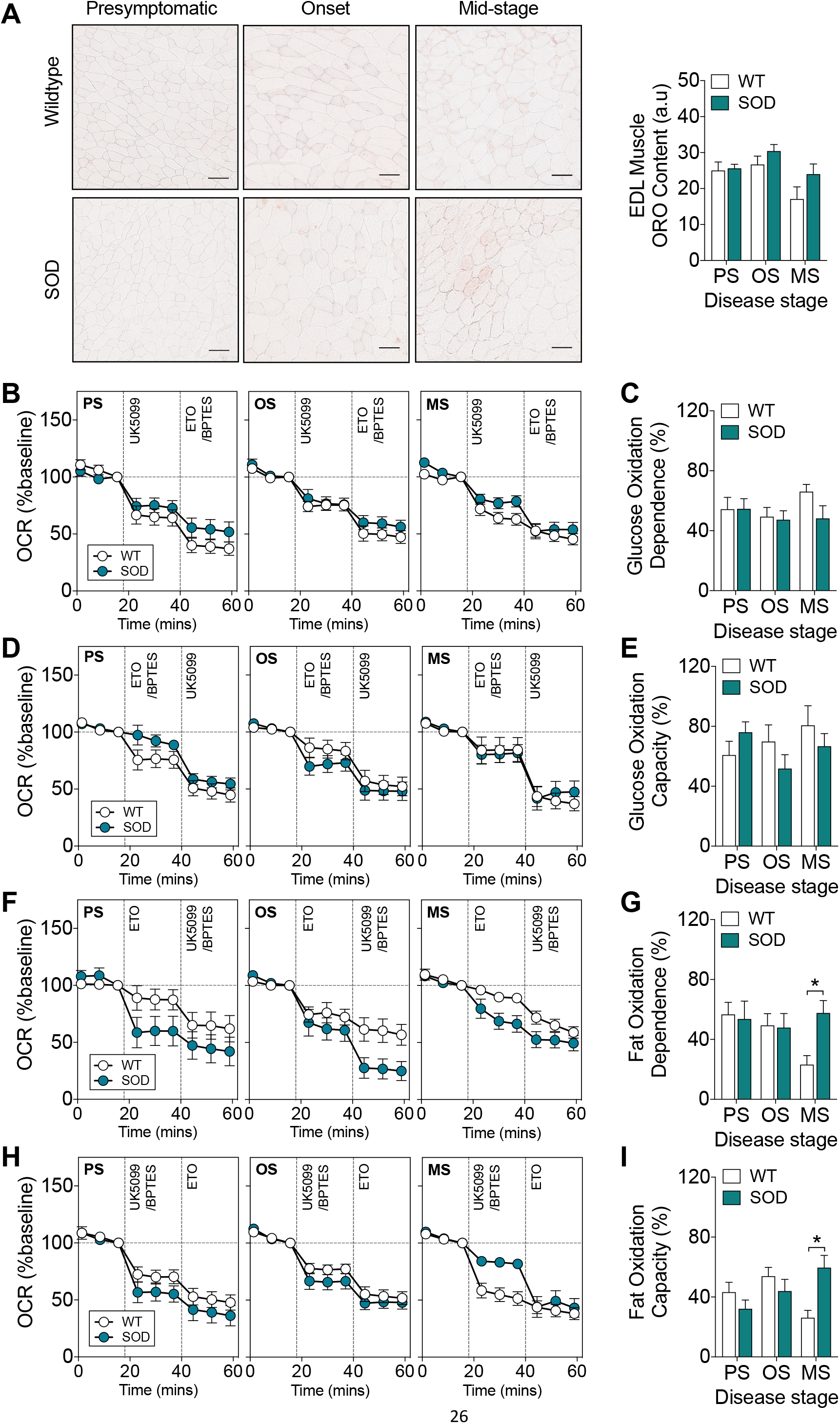
A functional shift in mitochondrial fuel preference from glucose to fat occurs in glycolytic extensor digitorum longus (EDL) muscle of SOD1^G93A^ mice. (A) Oil Red O staining and quantification of intramuscular lipid in the glycolytic EDL muscle of SOD1^G93A^ and age-matched wild-type (WT) mice (n=5/group), scale bar = 50μm. (B) Data trace of mitochondrial oxygen consumption rate (OCR, % baseline) contributed to by glucose oxidation dependence. (C) Quantification of glucose oxidation dependence in isolated EDL muscle fiber bundles from SOD1^G93A^ mice and WT controls. (D) Data trace of mitochondrial OCR (% baseline) contributed to by glucose oxidation capacity. (E) Quantification of glucose oxidation capacity in isolated EDL muscle fiber bundles from SOD1^G93A^ mice and WT controls. (F) Data trace of mitochondrial OCR (% baseline) contributed to by fat oxidation dependence. (G) Quantification of fat oxidation dependence in SOD1^G93A^ mice and WT control mice. (H) Data trace of mitochondrial OCR (% baseline) contributed to by fat oxidation capacity. (I) Quantification of fat oxidation capacity in SOD1^G93A^ mice and WT controls. White circles and columns represent WT mice; blue circles and columns represent SOD1^G93A^ transgenic mice. All data presented as mean ± SEM for n=5-12/group. \**P*<0.05, two-way ANOVA with Bonferroni’s post-doc test. PS, presymptomatic; OS, onset; MS, mid-stage.

To determine the capacity of glucose and fatty acid oxidation pathways to sustain mitochondrial respiration in the absence of substrate competition, and to study substrate utilization in the presence of increased energy demand, we measured the capacity of mitochondria in EDL fiber bundles to oxidize pyruvate or palmitate in the presence of the mitochondrial uncoupler, carbonyl cyanide-4-phenylhydrazone (FCCP; Figure 5A and B). The basal oxygen consumption rate (OCR) of EDL muscles from WT and SOD1^G93A^ mice was similar (Figure 5C). In the presence of pyruvate, we observed a significant elevation in maximal peak OCR in EDL muscle fiber bundles from SOD1^G93A^ mice at the onset stage of disease (Figure 5A). In the presence of palmitate, we observed no difference in maximal OCR in EDL muscle fiber bundles between SOD1^G93A^ and WT controls (Figure 5B). Compared to WT mice, total oxygen consumption in the EDL muscle fibers from SOD1^G93A^ mice was significantly higher at the mid-stage of disease when pyruvate was provided as the external energy substrate (Figure 5D). However, oxygen consumption between EDL muscle fibers from SOD1^G93A^ mice and WT mice was comparable when palmitate-BSA was provided as the external energy substrate (Figure 5E). Thus, despite exhibiting a functional preference towards fatty acid oxidation, glycolytic EDL muscle fibers from SOD1^G93A^ mice are capable of utilizing glucose metabolism pathways to sustain mitochondrial function when there are no competing fatty acid substrates.

**Figure 5:**
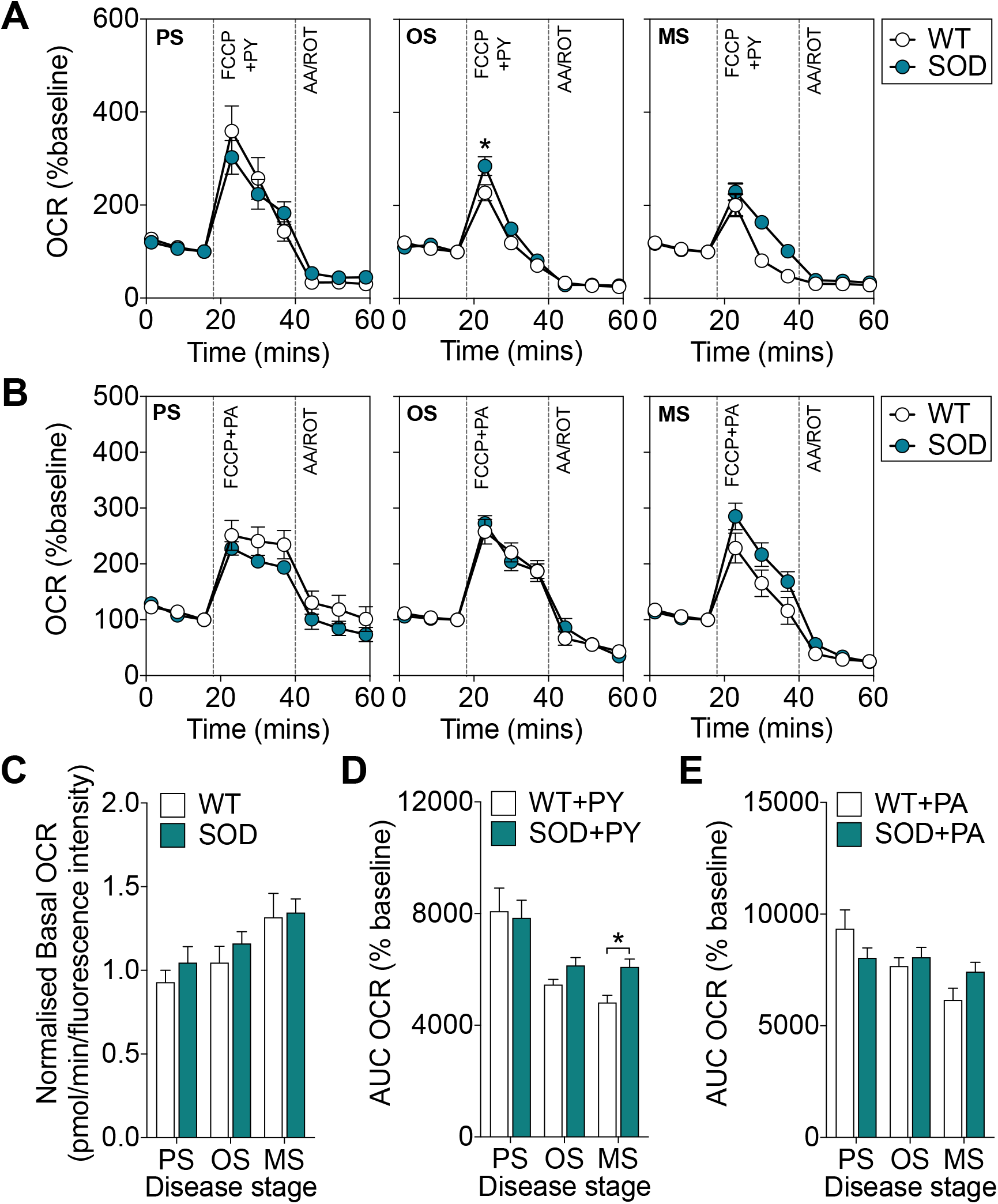
Glycolytic extensor digitorum longus (EDL) muscle fiber bundles from SOD1^G93A^ mice are capable of utilizing glucose metabolism pathways. (A) Data traces of mitochondrial oxygen consumption rate (OCR, % of baseline) when carbonylcyanide-p-trifluoromethoxyphenylhydrazone (FCCP) was used to induce maximal mitochondrial respiration in the presence of (A) pyruvate (PY) or (B) palmitate (PA). (C) Quantification of basal OCR in EDL muscle fiber bundles from SOD1^G93A^ mice and wild-type (WT) age-matched control mice. (D) Quantification of area under the curve (AUC) of OCR in muscle fiber bundles from SOD1^G93A^ and WT control mice in the presence of FCCP and pyruvate. (E) Quantification of area under the curve (AUC) of OCR in muscle fiber bundles from SOD1^G93A^ and WT controls in the presence of FCCP and palmitate. White circles and columns represent WT mice; blue circles and columns represent SOD1^G93A^ transgenic mice. All data presented as mean ± SEM for n=5-12/group. \**P*<0.05, two-way ANOVA with Bonferroni’s post-doc test. PS, presymptomatic; OS, onset; MS, mid-stage.

### ALS patient-derived myotubes have increased dependence on fat oxidation

We next aimed to determine whether mitochondrial fuel selection preference observed in SOD1^G93A^ mice was present in human ALS subjects. We generated primary myotubes from skeletal muscle biopsies obtained from ALS patients and age-matched healthy controls and conducted real-time assessment of substrate utilization. Demographics of our study population are detailed in Table 1. Sex, age, and clinical demographics of ALS patients are detailed in Supplementary Table 1. When compared to myotubes derived from healthy controls, myotubes from patients with ALS had similar levels of glucose oxidation dependency and capacity (Figure 6A). ALS patient derived myotubes also had similar levels of fat oxidation capacity when compared to myotubes derived from healthy controls. However, they exhibited an increased dependence on fat oxidation (Figure 6B).

**Figure 6:**
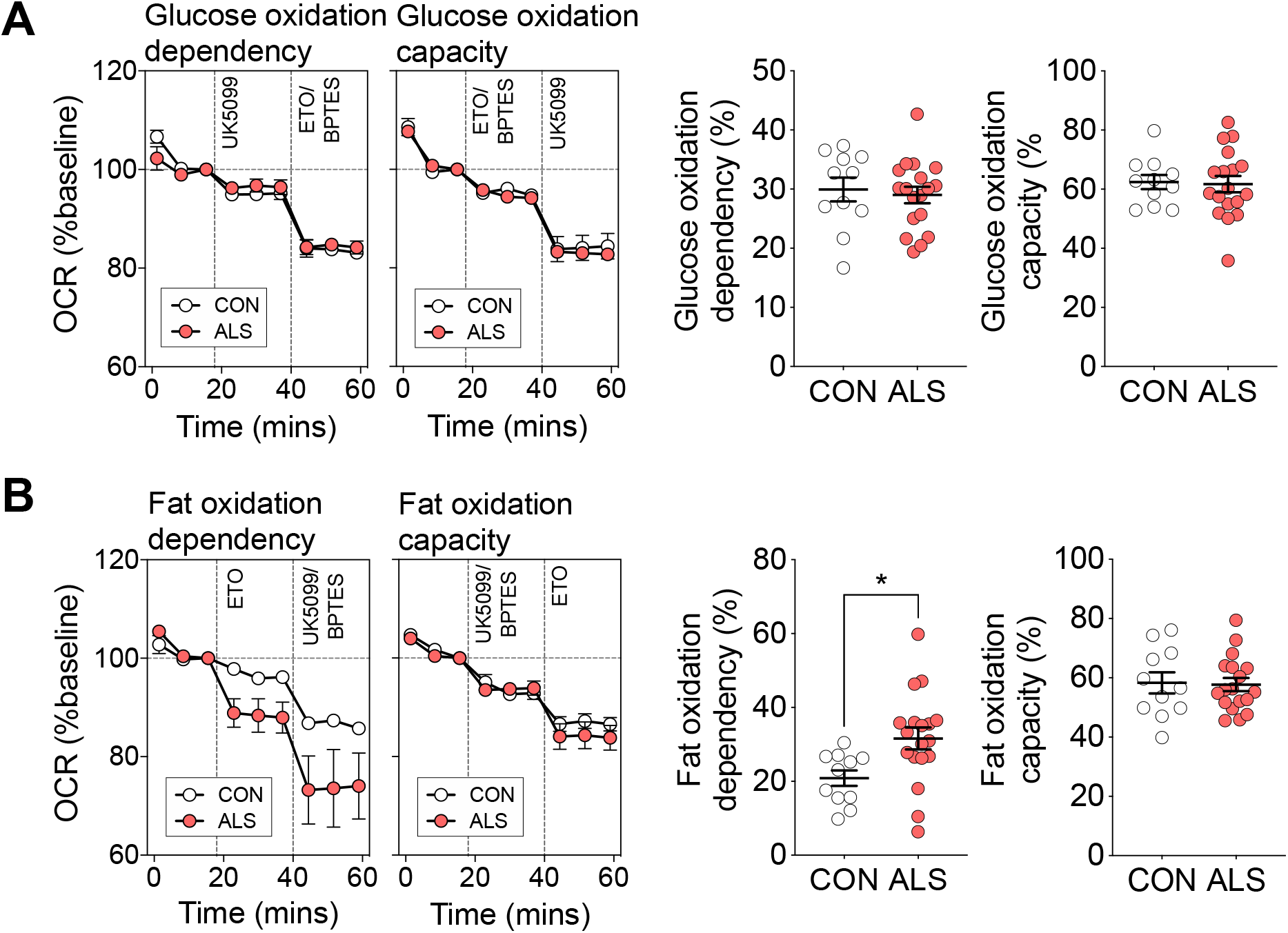
Myotubes derived from patients with amyotrophic lateral sclerosis (ALS) have increased dependence on fat oxidation. (A) Data traces, and quantification of mitochondrial oxygen consumption rate (OCR, % baseline) contributed to by glucose oxidation dependency and capacity in ALS and control myotubes. (B) Data traces, and quantification of mitochondrial OCR (% baseline) contributed to by fat oxidation dependency and capacity in ALS and control myotubes. All data presented as mean ± SEM for n=11 control and n=18 ALS individuals. \**P*<0.05, Mann Whitney t-test.

We next assessed the relationships between substrate oxidation in myotubes and the metabolic and clinical characteristics of our ALS cohort. The metabolic measures were resting energy expenditure and the metabolic index, which we have previously used to define hypermetabolism in ALS patients (Supplementary Table 2) (Steyn *et al*., 2018a). We found that glucose oxidation capacity was negatively correlated with resting energy expenditure, and that fat oxidation dependency was greater in myotubes derived from ALS patients with higher resting energy expenditure. Substrate oxidation was not correlated with the metabolic index.

We observed no relationship between substrate oxidation in ALS patient derived myotubes relative to the severity of disability as determined by the ALSFRS-R at the time of metabolic assessment (Supplementary Table 2). However, glucose oxidation capacity was greater, and fat oxidation dependence was lower in patients with a more rapidly progressing disease (indicated by a faster decline in ALSFRS-R scores (ΔFRS); Supplementary Table 2, Figure 7). Collectively, this data indicates that substrate utilization in myotubes is not related to hypermetabolism in ALS, but rather, it appears to be linked to the resting energy expenditure of ALS patients. Moreover, our data also suggest that substrate utilization in myotubes is associated with the rate of functional decline in patients with ALS.

**Figure 7:**
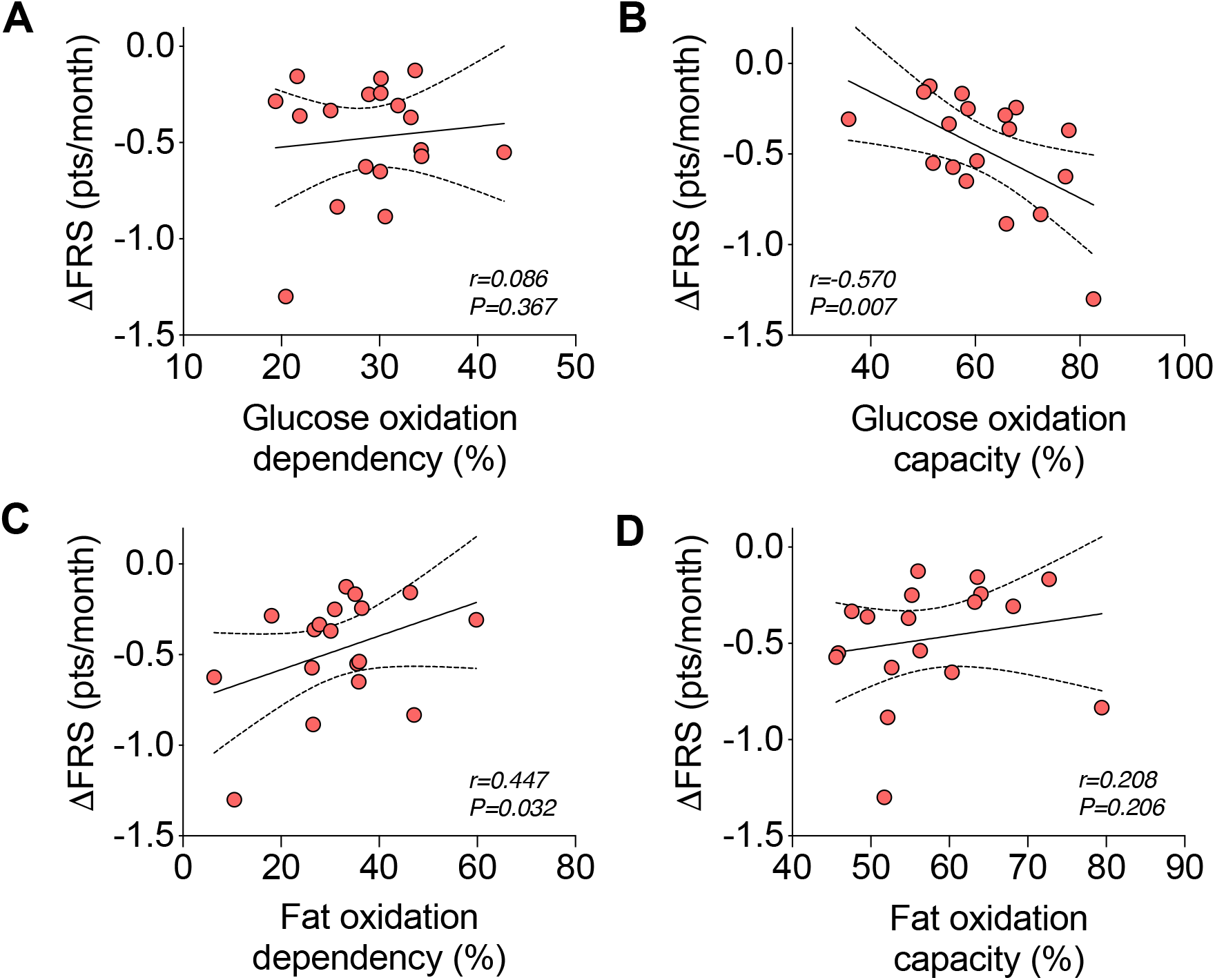
Resting energy expenditure and rate of functional decline in patients with amyotrophic lateral sclerosis (ALS) is linked to substrate utilization in myotubes derived from respective donors. Correlation analyses between rate of disease progression (ΔFRS) and (A) glucose oxidation dependency, (B) glucose oxidation capacity, (C) fat oxidation dependency, and (D) fat oxidation capacity. ΔFRS, rate of functional decline as defined by (amyotrophic lateral sclerosis function rating scale-revised score-48)/disease duration (in months) from symptom onset. Data presented as Pearson correlations for n=18 ALS patients.

## Discussion

The primary aim of this study was to explore potential mechanisms underpinning altered whole body metabolic balance in ALS (Ahmed *et al*., 2018; Steyn *et al*., 2018a; Vandoorne *et al*., 2018). We show that an increase in whole-body energy expenditure in symptomatic SOD1^G93A^ mice is associated with a decline in both fat mass and fat free mass. We also find that glycolytic muscle from symptomatic SOD1^G93A^ mice exhibit an increased dependence on fatty acids as an energy substrate, as well as an increased capacity to utilize fatty acids. In myotubes derived from patients with ALS, we show a similar association between dependence on fatty acid oxidation and resting energy expenditure. Further there was an association between muscle cellular metabolism and rate of disease progression.

Compared to WT littermates, SOD1^G93A^ mice exhibited an increase in total whole-body oxygen consumption (a proxy for energy expenditure) alongside a concomitant decline in body mass, fat mass and fat free mass over the course of disease. These observations are in line with the notion that increased energy expenditure in ALS contributes to weight loss (Dupuis *et al*., 2011). However, given that hypermetabolism has not been shown to be associated with weight loss in human ALS (Steyn *et al*., 2018b), these divergent observations between ALS mice and patients highlight that multiple factors contribute to weight loss is patients with ALS. In patients with ALS, weight loss is likely to occur due to a combination of factors, including reduced capacity to meet energy requirements (Ngo *et al*., 2017) (including loss of appetite resulting in loss of fat mass (Ngo *et al*., 2019; Mezoian *et al*., 2020)), as well as neurogenic wasting (Al-Sarraj *et al*., 2014).

A failure to accumulate fat mass was an early feature of disease in SOD1^G93A^ mice, and this is associated with sustained lipolysis rather than reduced food intake. Why high levels of lipolysis is maintained throughout disease course in SOD1^G93A^ mice remains to be determined. It is well established that lipolysis and fatty acid mobilization are upregulated in response to increased muscle energy requirements, for example, during exercise (Goodpaster and Sparks, 2017). Thus, it has been proposed that increased mobilization of lipids in ALS mice could sustain metabolic requirements in peripheral skeletal muscle (Dupuis *et al*., 2004; Fergani *et al*., 2007). Expanding on previous reports of increased expression of genes associated with fatty acid oxidation in skeletal muscle of SOD1^G86R^ mice (Palamiuc *et al*., 2015), we have now generated the first evidence to show that there is a greater functional dependence and capacity of glycolytic skeletal muscle from symptomatic SOD1^G93A^ mice to utilize fatty acids as a fuel substrate. This increased dependence and capacity for fatty acid oxidation could be an adaptive response of the muscle due to reduced muscle glucose uptake and glucose intolerance (Pradat *et al*., 2010; Desseille *et al*., 2017). Yet, we found that glycolytic muscle fibers were able to utilize pyruvate, a product of glucose oxidation, following inhibition of fatty acid and glutamine pathways. While providing evidence to suggest that glucose oxidation mechanisms remain intact in glycolytic SOD1^G93A^ muscle, our data lend further support to the notion that increased utilization of fatty acids could inhibit the use of glucose in ALS (Palamiuc *et al*., 2015), presumably via the Randle cycle (Hue and Taegtmeyer, 2009).

In order to understand whether the metabolic changes observed in isolated SOD1^G93A^ mouse muscle fibers occur in human ALS, we investigated substrate oxidation capacity and dependence in myotubes derived from patients with ALS. Similar to SOD1^G93A^ mouse muscle, we observed an increase in fatty acid oxidation dependence in ALS patient myotubes, although fatty acid oxidation capacity remained unchanged. In addition, we found that myotubes with higher fat oxidation dependence were derived from ALS patients with higher resting energy expenditure. These results are congruent with the known ability for skeletal muscle to adapt to changes in energy supply and requirement to maintain energy homeostasis (Horowitz, 2003). It is well known that fatty acid metabolism provides more ATP when compared to glucose metabolism (Turner *et al*., 2014). As such, an increase in the need for fatty acid oxidation in myotubes obtained from patients with higher resting energy expenditure may function to increase ATP availability. Moreover, the correlation observed between resting energy expenditure and fatty acid oxidation dependence aligns with physiological processes wherein continued dependence on fatty acid metabolism requires greater levels of oxygen consumption when compared to glucose metabolism (Turner *et al*., 2014).

We did not observe any relationship between the metabolic index or any substrate oxidation parameters in ALS patient myotubes. We assume that the findings in myotubes reflect the situation in intact muscle. Hypermetabolism in ALS could arise from other tissues such as the brain. Previous investigation using [^18^F]fluorodeoxyglucose positron emission tomography supports such a hypothesis, as increased energy use has been observed across a number of brain regions in ALS patients (Cistaro *et al*., 2012). In those ALS patient who show hypermetabolism, the metabolic changes could reflect a whole-body response to disease. Indeed, hypermetabolism is a feature of critical illness, wherein the degree of hypermetabolism varies with the severity and duration of illness (McClave and Snider, 1994).

We found that myotubes with higher fatty acid oxidation dependence were derived from patients with slower clinical decline, indicating that substrate availability and/or use could be a factor that determines disease progression in ALS. Previous studies linking higher levels of serum lipids and fat mass with longer survival (Marin *et al*., 2011; Lindauer *et al*., 2013; Huang *et al*., 2015) provide some evidence to suggest that lipids are beneficial in ALS.

Although we have used both mouse and human-derived models to investigate aspects of muscle metabolism in ALS, there are some limitations to our study. First, the SOD1^G93A^ mouse, albeit an accepted pre-clinical model of ALS, is representative of only a small proportion of ALS patients. However we found similar alterations in substrate oxidation in the myotubes derived from some ALS patients, indicating that muscle metabolic changes are possibly present widely in the disease. Second, while we conducted assessment of energy expenditure in ALS patients using indirect calorimetry (Haugen *et al*., 2007), and assessed substrate oxidation in real-time using Seahorse technology, our use of primary human myotubes is a caveat. Previously, it has been shown that human primary myotubes mirror the metabolic phenotypes of their donors (Ukropcova *et al*., 2005). Regardless, as primary myotubes are grown from cells which have been isolated from muscle biopsies that were removed from their physiological milieu, future studies assessing substrate oxidation in muscle *in vivo* are needed.

In summary, we demonstrate that an increase in fatty acid oxidation dependence and capacity occurs in glycolytic muscle of SOD1^G93A^ mice during the symptomatic stages of disease when they exhibit increased energy expenditure. In myotubes derived from ALS patients, a similar increase in fatty acid oxidation dependence occurs. While this change in fatty acid oxidation appears be associated with the progression of disease, it is not linked to hypermetabolism in human ALS. Given the heterogeneity of disease in ALS, there remains an important need for further studies that delineate mechanisms of metabolic imbalance and the link between substrate utilization and energy expenditure throughout the course of the disease. A comprehensive understanding of specific metabolic changes at an individual patient level will be essential for the development of treatments that aim to target metabolic pathways in ALS.

## Author contributions

FJS, TYX, DK, FG, SC, and STN conducted the experiments. TJT and SEK generated and maintained human myoblasts. EW and LR collected skeletal muscle biopsies. JSC provided infrastructure and materials. FJS and STN recruited ALS patients. PAM and RDH confirmed diagnoses of ALS patients. FJS, SEK, TWT, TYX, SC, WML, AF, CV, FR, JPL, and STN analyzed and interpreted the data. FJS and STN conceived and designed the study. FJS, SEK, TWT, and STN wrote and conducted the final review of the manuscript. All authors revised the manuscript.

## Acknowledgements

The authors gratefully acknowledge the assistance and support of staff at the University of Queensland Biological Resources (UQBR), and The Centre for Integrated Physiology at the School of Biomedical Sciences, University of Queensland. This work was funded by grants from The University of Queensland (NSRFS 2012002126 to STN), the Motor Neurone Disease Research Institute of Australia (Grant-in-aid, Graham Smith MND Research Grant (to FJS, PAM, and STN) and Charcot MND Research Grant to FJS, JSC, PAM, RDH and STN), the National Health and Medical Research Council (1101085 to FJS and STN, 1121962 to FCG), and AriSLA Foundation and AFM-Telethon (HyperALS and AFM-Telethon #22509 to AF, JPL, FR and CV). TYX was supported by international postgraduate research scholarships from the Australian Government, and University of Queensland centennial scholarships. STN was funded by the Scott Sullivan MND Research Fellowship (Queensland Brain Institute, the Royal Brisbane & Women’s Hospital Foundation, The MND and Me Foundation), and the Australian Institute for Bioengineering and Nanotechnology.

---

**Supplementary Table 1:**
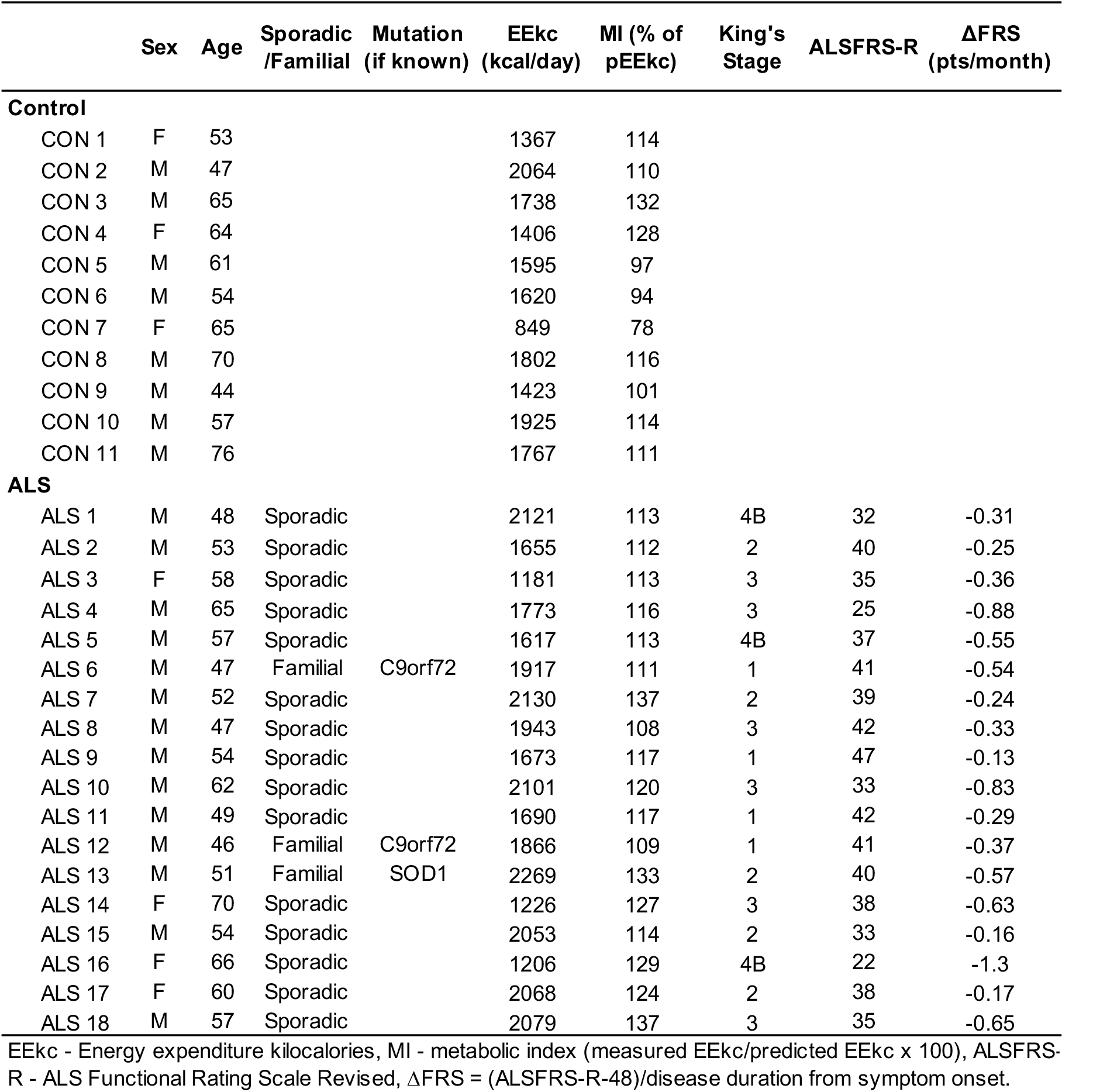
Sex and age for study participants. For ALS patients, metabolic and clinical demographics are included.

**Supplementary Table 2:** Correlations between cellular bioenergetic parameters and patient resting energy expenditure, metabolic index and clinical demographics.

|  | r | 95% confidence interval | P value |
| --- | --- | --- | --- |
| **Resting energy expenditure (mEEkc)** |  |  |  |
| Glucose oxidation dependence | 0.281 | -0.214 to 0.661 | 0.129 |
| Glucose oxidation capacity | -0.486 | -0.777 to -0.025 | **0.020** |
| Fat oxidation dependence | 0.685 | 0.320 to 0.873 | **0.001** |
| Fat oxidation capacity | 0.458 | -0.011 to 0.762 | **0.028** |
| **Metabolic Index (mEEkc/pEEkc)** |  |  |  |
| Glucose oxidation dependence | -0.030 | '-0.4898 to 0.4433" | 0.454 |
| Glucose oxidation capacity | 0.252 | '-0.2435 to 0.6432" | 0.157 |
| Fat oxidation dependence | -0.228 | '-0.6277 to 0.2678" | 0.182 |
| Fat oxidation capacity | 0.130 | '-0.3585 to 0.5628" | 0.303 |
| **ALSFRS-R** |  |  |  |
| Glucose oxidation dependence | 0.300 | -0.194 to 0.673 | 0.113 |
| Glucose oxidation capacity | -0.246 | -0.639 to 0.250 | 0.163 |
| Fat oxidation dependence | 0.021 | -0.450 to 0.483 | 0.466 |
| Fat oxidation capacity | -0.075 | -0.524 to 0.406 | 0.383 |
| **ΔFRS (pts/month)** |  |  |  |
| Glucose oxidation dependence | 0.086 | -0.397 to 0.531 | 0.367 |
| Glucose oxidation capacity | -0.570 | -0.819 to -0.141 | **0.007** |
| Fat oxidation dependence | 0.447 | -0.026 to 0.756 | **0.032** |
| Fat oxidation capacity | 0.208 | -0.287 to 0.615 | 0.206 |

mEEkc - measured energy expenditure as kilocalories/day, pEEkc - predicted energy expenditure as kilocalories/day, ALSFRS-R - ALS Functional Rating Scale Revised, ΔFRS = (ALSFRS-R-48)/disease duration (in months) from symptom onset.

